# Structural characterization and evolutionary analyses of the *Coccidioides immitis* and *Coccidioides posadasii* mitochondrial genomes

**DOI:** 10.1101/2020.09.14.296954

**Authors:** Marcus de Melo Teixeira, B. Franz Lang, Daniel R. Matute, Jason E. Stajich, Bridget Barker

## Abstract

Fungal mitochondrial genomes encode for genes involved in crucial cellular processes, such as oxidative phosphorylation and mitochondrial translation, and these genes have been used as molecular markers for population genetics studies. *Coccidioides immitis* and *C. posadasii* are endemic fungal pathogens that cause coccidioidomycosis in arid regions across both American continents. To date, almost one hundred *Coccidioides* strains have been sequenced. The focus of these studies has been exclusively to infer patterns of variation of nuclear genomes (nucDNA). However, their mitochondrial genomes (mtDNA) have not been studied. In this report, we describe the assembly and annotation of mitochondrial reference genomes for two representative strains of *C. posadasii* and *C. immitis*, as well as assess population variation among 77 published genomes. The circular-mapping mtDNA molecules are 68.2 Kb in *C. immitis* and 75.1 Kb in *C. posadasii*. We identified the fourteen mitochondrial protein-coding genes common to most fungal mitochondria, including genes encoding the small and large ribosomal RNAs (*rns* and *rnl*), the RNA subunit of RNAse P (*rnp*B), and 26 tRNAs organized in polycistronic transcription units, which are mostly syntenic across different populations and species of *Coccidioides*. Both *Coccidioides* species are characterized by a large number of group I and II introns, harboring twice the number of elements as compared to closely related Onygenales. The introns contain complete or truncated ORFs with high similarity to homing endonucleases of the LAGLIDADG and GIY-YIG families. Phylogenetic comparison of the mtDNA and nucDNA genomes shows discordance, possibly due to differences in patterns of inheritance. In summary, this work represents the first complete assessment of mitochondrial genomes among several isolates of both species of *Coccidioides*, and provides a foundation for future functional work.

## Introduction

Fungal mitochondrial genomes exist as either linear or circular-mapping molecules and range from ~17.6 kb (e.g. *Schizosaccharomyces pombe* Genbank ID MK618090.1) to well over 200 kb (e.g. 272,238 bp in *Morchella importuna* (1)). Fungal mitochondrial genomes usually encode proteins involved in oxidative phosphorylation - the main source of ATP production of the cell - as well as two ribosomal RNA subunits, and a set of tRNAs involved in mitochondrial ribosome translation. More specifically, fungal mitochondrial protein-coding genes fall into several classes: seven subunits of ubiquinone oxidoreductase (*nad;* not present in a number of Saccharomycotina and in fission yeasts, (2)), cytochrome b (*cob*), three subunits of cytochrome oxidase (*cox*) and up to three ATP synthase subunits (*atp*; the presence of *atp8* and *atp9* varies among fungal taxa) (3). Also, a gene encoding a ribosomal protein subunit (*rps3*) is present in most fungal mitochondrial genomes. Mitochondrial protein-coding genes are frequently intercalated with genes that encode structural RNAs: ribosomal RNAs (small and large subunit rRNAs *rns* and *rnl*), the RNA subunit of RNase P (*rnpB*) with infrequent occurrence across fungi, and variable numbers of tRNAs. Notable exceptions are the *nad* genes, which tend to be organized in operon-like structures, with some of the genes overlapping without discernable intergenic regions (e.g., *nad4L* situated upstream of *nad5*, overlapping by one to a dozen or more nucleotides) (3).

Mitochondrial genes in fungi contain highly variable numbers of group I and II introns that are inserted in protein-coding as well as rRNA genes (4). For instance, *Endoconidiophora* species seem to contain more than 80 mitochondrial introns (5), which can create gene annotation challenges especially when transcriptome data are not available. Both intron groups may contain complete or truncated ORFs that encode either homing endonucleases of the LAGLIDADG and GIY-YIG families, or reverse transcriptases/maturases (6). If present, these proteins direct an intron transfer within mitochondrial genomes of genetically compatible fungal isolates, or less frequently across genera, and even kingdom boundaries (7). Mitochondrial DNA (mtDNA)-encoded genes are particularly prone to crossing species boundaries. As intron transfer *via* homing endonucleases involves genetic co-conversion of flanking exon sequences, phylogenetic inferences using mtDNA— especially genes with high intron numbers (e.g., *cox1, cob* and *rnl* (3, 8))— may reveal replacement of coding regions, related to ongoing intron invasion.

In this study, we focus on describing the mitogenomes of *Coccidioides immitis* and *C. posadasii* (Ascomycota, Onygenales), which are fungal species endemic to both American continents, and the causative agents of coccidioidomycosis (9). This disease is most frequently reported in the “Lower Sonoran Life Zone” in California, Arizona, Texas, and northwestern Mexico (10). However, the disease is also reported in arid and semi-arid areas throughout the American continents (11). The two species have a complex evolutionary history dominated by biogeographic distribution patterns (12, 13). *Coccidioides immitis* has been found in California and Baja Mexico as well in eastern Washington state, and each region harbors unique genotypes (14-16). *Coccidioides posadasii* is present throughout Arizona, Texas, Central, and South America, and population structure has been described as containing an Arizona population, a Texas/Mexico/South America (TX/MX/SA) population, and a distinct Caribbean population (13).

Notably, nucDNA studies have found extensive differentiation between species of *Coccidioides* with some evidence for gene flow between species (17, 18). The two species, *C. immitis* and *C. posadasii*, can be discriminated based on polymorphisms found at the first intron of the *cox1* gene (19). Yet, no studies have addressed whether or not mtDNA reflects the divergence of ncDNA, or if mtDNA has moved between *Coccidioides* species or among populations. In this study we: *i*) describe the full circular-mapping mitogenomes of *C. posadasii* and *C. immitis, ii*) compare their core genes, structural RNAs and introns of group I and II with other Onygenales fungal species, and *iii*) compare the evolutionary trajectories between the mtDNA and nucDNA genomes among publicly available genomes of this medically important fungal pathogen.

## Materials and Methods

### Mitochondrial genome assembly and annotation

Paired end Illumina sequence reads from 20 *Coccidioides immitis* and 57 *C. posadasii* were retrieved from the Sequence Read Archive (SRA) and accessions and details are listed in **Table S1**. Following cleaning and quality-clipping of reads with Trimmomatic v0.35, we assembled the genomes of *C. posadasii* Tucson-2 and *C. immitis* WA221 using the SPAdes Genome Assembler v3.14.0 (20) with a kmer sizes 61, 91, and 127. We identified mitochondrial contigs in this initial assembly using similarity searches with expected fungal genes. To minimize assembly error we (i) used Rcorrector [Song, L., Florea, L. Rcorrector: efficient and accurate error correction for Illumina RNA-seq reads. GigaSci 4, 48 (2015).] for read correction, (ii) reduced the number of Illumina reads to a target kmer coverage of the mtDNA between 30-50x, (iii) reads mapping against the identified mitochondrial contigs were identified with Bowtie2 (21), which were then (iii) reassembled with Spades, resulting in preliminary (uncorrected) mitogenome assemblies. In a final step, all reads of the reduced 30-50x read set were aligned back to the preliminary assembly with Bowtie2 and analyzed for kmer coverage with Bedtools v2.29.2 (22). We identified incorrectly-assembled reads, defined by kmer frequency values of two or lower (likely the result of hybrid reads, originating from ligation of unrelated genomic DNA fragments during library construction), and removed them from the final assemblies. For both species, we obtained single circular-mapping closed contigs that carry the expected full set of fungal mitochondrial genes.

To compare the *Coccidioides* mtDNA assembles with other fungi, we retrieved full mitochondrial assemblies from other Onygenales available in the NCBI GenBank database: *Histoplasma capsulatum* H143 (GG692467.1, direct submission) *Paracoccidioides brasiliensis* Pb18 (AY955840.1, (23)), *Blastomyces dermatitidis* ATCC 18188 (GG753566.1, direct submission), *Epidermophyton flocossum* ATCC 26072 (AY916130.1, (24)), *Trichophyton rubrum* BMU 01672 (FJ385026.1, (25)) and *Ascosphaera apis* ARSEF 7405 (AZGZ01000045.1, (26)). Other close relatives of *Coccidioides* (e.g. *Uncinocarpus*) had incomplete mitogenomes (27).

Mitochondrial genes as well as introns of group I and group II, tRNAs, RNase P RNA (*rnpB*), and the small and large subunit rRNAs (*rns* and *rnl*) for *Coccidioides* and other related Onygenalean fungi were annotated using the MFannot pipeline (https://megasun.bch.umontreal.ca/cgi-bin/dev_mfa/mfannotInterface.pl; https://github.com/BFL-lab/Mfannot). *Coccidioides* annotations were manually inspected and intron boundaries were checked and adjusted by aligning available RNAseq data (27) with respective mitochondrial assemblies using Bowtie 2 (21). The assemblies and annotations were deposited in GenBank (accession numbers TBD) and were visualized with the OGDRAW pipeline (28).

### Single nucleotide polymorphism assessment and phylogenetic analysis

SNPs from 77 *Coccidioides* isolates were identified among the mitochondrial genomes. We mapped Illumina paired-end reads into individual mitochondrial coding-genes using Burrows-Wheeler Aligner (BWA) v 0.7.7 (29) to assembled mitochondrial references *C. posadasii* strain Tucson2 or *C. immitis* strain WA221. Indels were realigned to its reference genomes using GATK RealignerTargetCreator and IndelRealigner tools (GATK toolkit v 3.3-0 (30)). To call SNPs, we used the UnifiedGenotyper package. We only included SNPs not located in potentially duplicated loci (as identified by NUCmer, (31)), with more than 10X coverage, and with a minor allele frequency of at least 10%. We used the same approach to call SNPs for the nucDNA genomes (13). We generated Maximum Likelihood (ML) concatenated trees for mtDNA and nucDNA using methods implemented in IQ-TREE software (32) using -m MFP option (ModelFinder - (33)) for model selection and 1,000 ultrafast bootstraps coupled with Shimodaira–Hasegawa-like approximate likelihood ratio test (SH-aLRT) were performed for branch confidence test (34).Finally, we compared the topology of the two trees using FigTree v1.4.2 - http://tree.bio.ed.ac.uk/software/figtree/, and scored the disagreements the two topologies were using TOPD/FMTS v 4.6 (35).

## Results

### The *Coccidioides* spp. mitogenome

We assembled complete circular mtDNA molecules for each of the two species of *Coccidioides*. The two assemblies differ in size: the mtDNA genome is 68.6 Kb in *C. immitis* and 75.1 Kb in *C. posadasii* (Figure 1). There is variation in mtDNA genome size among Onygenales (Table 1). The mtDNA of both species of *Coccidioides* are on the larger end of the continuum. The mitogenomes of *Coccidioides* harbor 14 protein-coding genes responsible for the formation of ubiquinone oxidoreductase, cytochrome b, cytochrome oxidase and ATP synthase protein complexes (Figure 1, Figure 2). The two ribosomal small and large subunit rRNA genes (*rns* and *rnl*), RNase P RNA (*rnpB*) and 26 tRNAs organized in polycistronic transcription units are all present. The gene composition and synteny are conserved between *Coccidioides* (Onygenaceae), *Blastomyces, Histoplasma*, and *Paracoccidioides* (all Ajellomycetaceae) (Figure 1, (36)). The position of the gene *atp8* differs between the Onygenaceae/Ajellomycetaceae species and other species of the Onygenales, such as dermatophytes (*Trichophyton rubrum and Epidermophyton floccosum -* Arthrodermataceae), and the bee-pathogenic fungus *Ascosphaera apis* (Ascosphaeraceae, Figure 2). We observed no gene gain and losses of core mitochondrial genes within Onygenalean fungi (Figure 2).

**Table 1.** Numbers of introns and classes among Onygenalean fungi

|  | *C.*<br>*immitis* | *C.*<br>*posadasii* | *H.*<br>*capsulatum* | *P.*<br>*brasiliensis* | *B.*<br>*dermatitidis* | *E.*<br>*flocossum* | *T.*<br>*rubrum* | *A. apis* |
| --- | --- | --- | --- | --- | --- | --- | --- | --- |
| **Mitochondrial**<br>**genome size (bp)** | 68,597 | 75,194 | 39,129 | 71,335 | 51,071 | 30,910 | 26,985 | 118,650 |
| **Intron IB**<br>**(complete)** | 15 | 15 | 3 | 5 | 6 | 3 | 0 | 8 |
| **intron IB (extra**<br>**insertion)** | 0 | 0 | 1 | 0 | 0 | 0 | 0 | 0 |
| **intron IB (5',**<br>**partial)** | 0 | 1 | 0 | 0 | 0 | 0 | 0 | 1 |
| **intron IB (3',**<br>**partial)** | 1 | 2 | 1 | 0 | 2 | 0 | 0 | 1 |
| **intron IA** | 3 | 3 | 1 | 0 | 1 | 1 | 2 | 1 |
| **intron IA (5',**<br>**partial)** | 0 | 1 | 0 | 0 | 0 | 0 | 0 | 1 |
| **intron I (derived,**<br>**A)** | 1 | 1 | 0 | 0 | 2 | 1 | 0 | 5 |
| **intron I (derived,**<br>**B1)** | 2 | 2 | 0 | 4 | 0 | 0 | 0 | 11 |
| **intron ID** | 4 | 3 | 1 | 0 | 1 | 1 | 0 | 5 |
| intron IC1 | 0 | 0 | 0 | 0 | 0 | 0 | 0 | 2 |
| intron IC2 | 4 | 4 | 0 | 1 | 1 | 0 | 0 | 5 |
| intron I (derived, B2) | 1 | 1 | 1 | 0 | 1 | 0 | 0 | 1 |
| intron II (domain V) | 4 | 5 | 1 | 3 | 2 | 0 | 0 | 3 |
| intron II, derived | 1 | 1 | 0 | 0 | 0 | 0 | 0 | 0 |
| Total | 36 | 39 | 9 | 13 | 16 | 6 | 2 | 44 |

**Figure 1.**
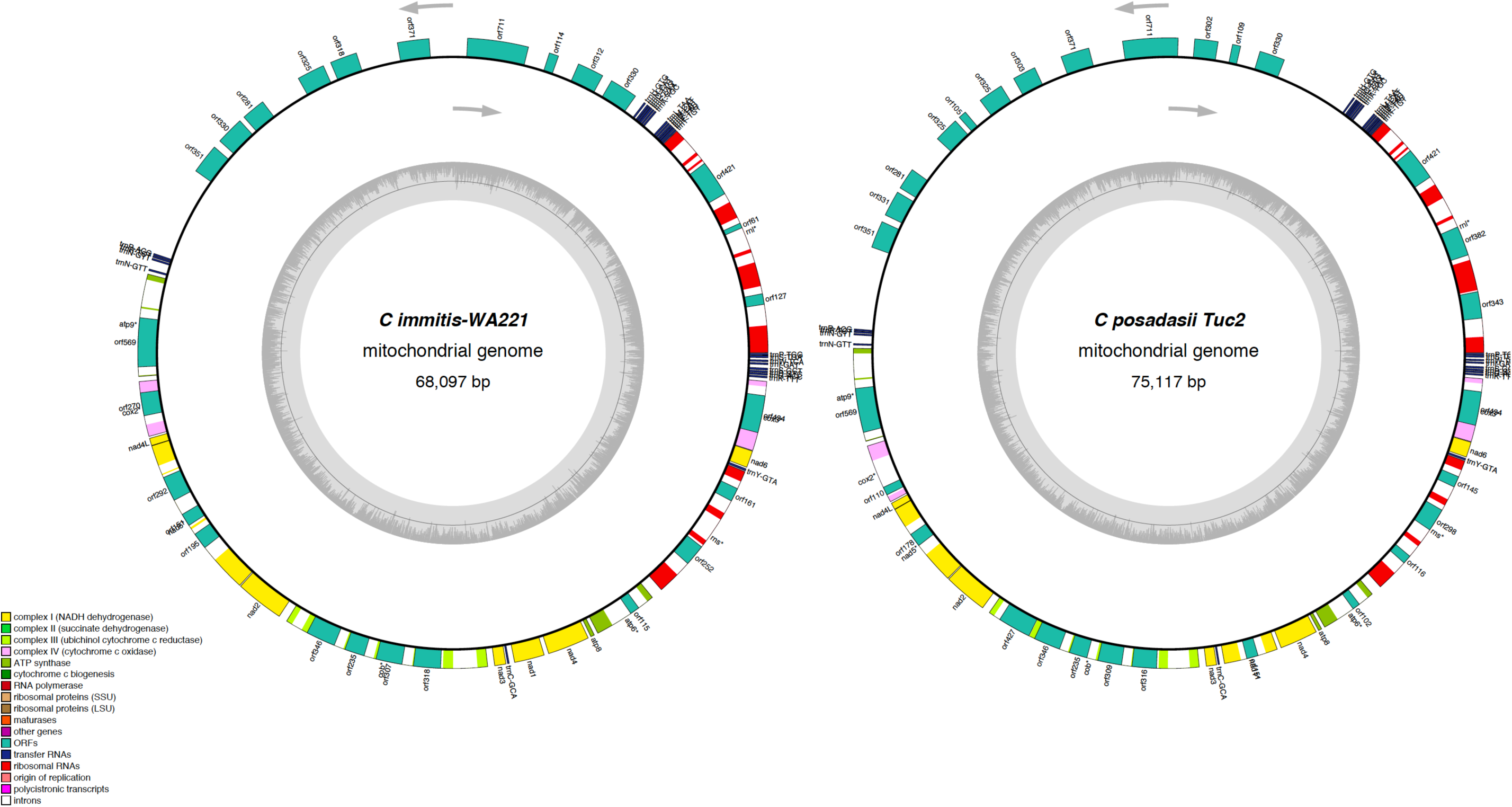
Circular maps of *C. immitis* and *C. posadasii* mitogenomes. The assembled and annotated genome features were converted into Genebank format and loaded into the OGDraw pipeline for physical visualization of the coding and non-coding elements of the mitochondrial genomes.

**Figure 2.**
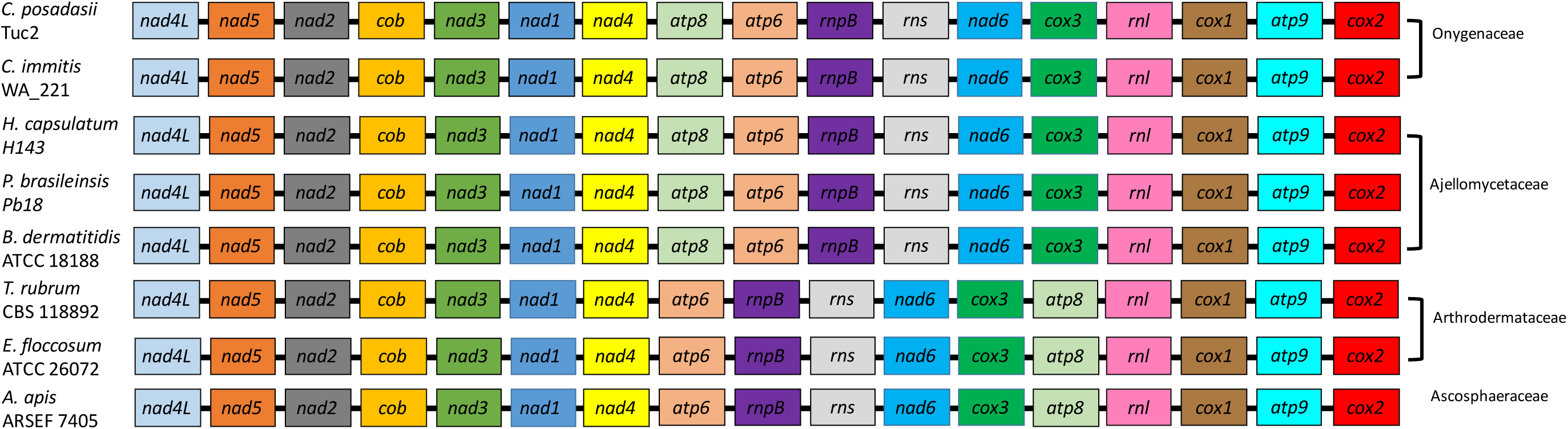
Mitochondrial gene content and synteny among Onygenalean fungi. Genes as color-coded (see legend) and displayed according positioning on the genome. The mitogenomes genomes are highly syntenic but *atp8* gene positioning is divergent between those fungal families.

The large size of the mtDNA genome in *Coccidioides* is due to the presence of introns and intron-encoded open reading frames (ORFs) in both *Coccidioides* species, resulting in a dramatic increase of intron type I and intron type II (Table 1, Figure 1) compared to other Ajellomycetaceae fungi. In fact, *Coccidioides* harbors twice the number of elements found in *B. dermatitidis*. The dermatophyte genera, *Epidermophyton* and *Trichophyton*, have only six and two intron elements respectively, whereas *C. immitis* and *C. posadasii* contain 39 elements respectively (Table 1). The introns found in the *Coccidioides* mitogenomes contain complete or truncated ORFs with high similarity to homing endonucleases of the LAGLIDADG and GIY-YIG families (Table 1). Both species contain 15 complete copies of Intron IB (Table 1). *Ascopharaceae apis* also has a large mtDNA genome (118.65Kb) and a high number of intron-type I, specifically the intron I - derived, B1 element (Table 1). The frequency and distribution of intron-types I and II in the genes *nadh5, cob* and *cox1* differ between *C. immitis* and *C. posadasii*, but these features are not differentiated within the species-complexes (Figure 1, Figure 2).

### Comparing mtDNA and nucDNA whole-genome trees

Finally, we compared the mitochondrial phylogeny with the whole genome species phylogeny. To score the differences in partitions produced between mtDNA and nucDNA trees, we used TOPD (35). If two trees are completely congruent the split distance score is 0, whereas with complete tree disagreement the score is 1. Differences are due to dissimilarity between the topologies as well as the number of overlapping taxa. The split distance is the ratio of (different/possible) between the *Coccidioides* mtDNA and nucDNA phylogenetic tree is 0.89 (132 /148), thus indicating that overall topologies are consistent. Both topologies support species divergence between *C. immitis* and *C. posadasii* (Figure 3). The mtDNA tree topology, shows a clear distinction between *C. immitis* and *C. posadasii* with no isolates being assigned to a different species. This results suggest no mtDNA gene exchange.

**Figure 3.**
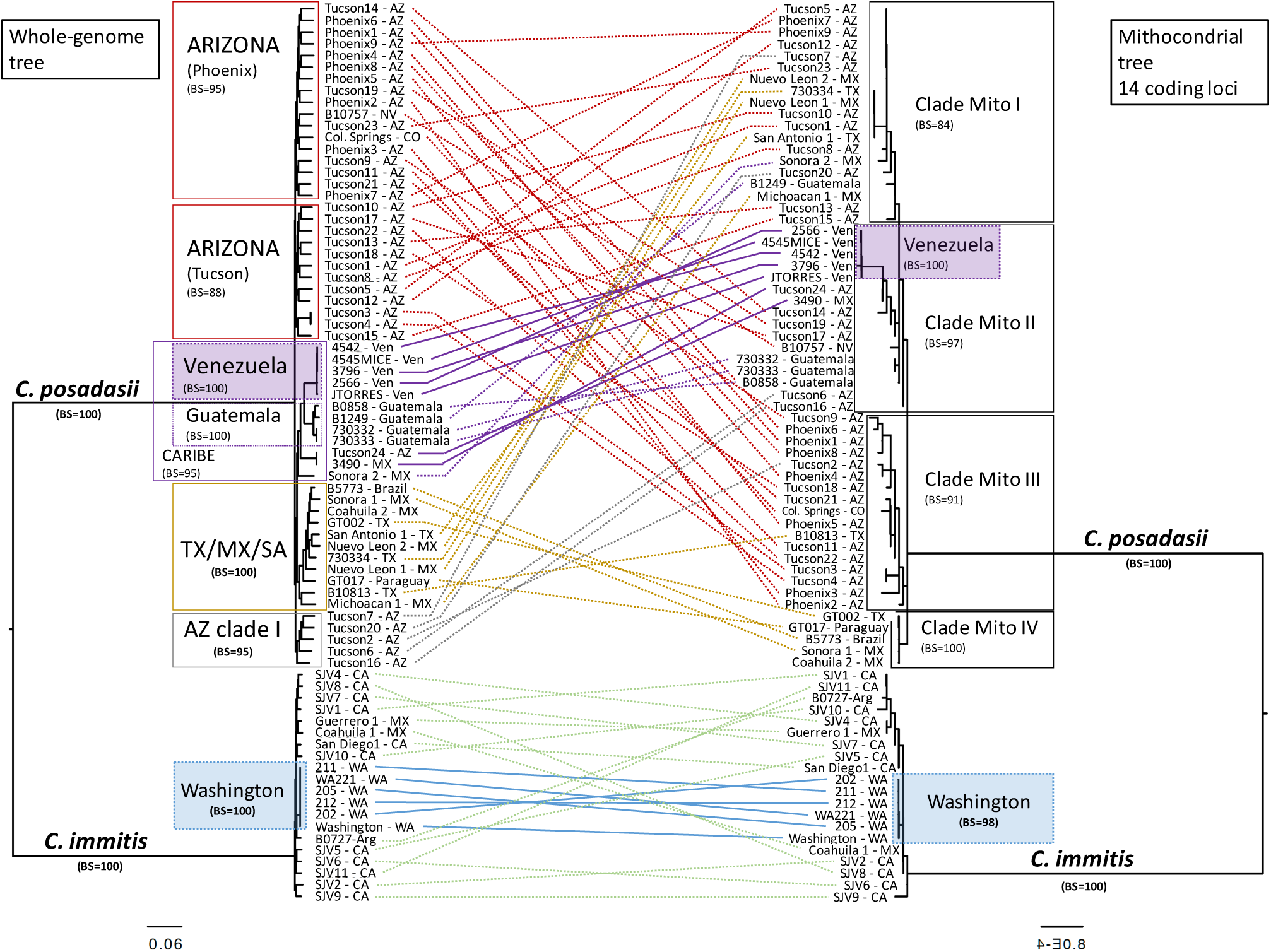
Tree topology comparisons of *Coccidioides* nucDNA (left panel) and mtDNA (right panel) phylogenomic trees. Phylogenetic tree branches are proportional to the nucleotide divergence (see scale) and the main clades are highlighted. Bootstrap support was calculated, and branch support was added to the corresponding clade. The terminal taxa are color-coded according to their placement on the nucDNA tree and taxa are connected between mtDNA and nucDNA phylogenomic trees in order to visualize concordance (solid lines) vs discordance (dotted lines).

Those populations have low intraspecific genetic variation, suggesting either strict clonal dispersion, or a recent founder effect (13).

Even though the main shape of the two topologies were consistent with each other. We also observed differences. Consistent with previous results, the nucDNA tree topology revealed three *C. posadasii* monophyletic populations: Arizona, Mexico/Texas/South America (MX/TX/SA) and Caribbean (13). Based on mtDNA among the three main *C. posadasii* populations that have been previously defined by markers differ substantially (Figure 3). Specifically, the mtDNA phylogeny shows *C. posadasii Arizona, Texas/Mexico/South America*, and *Caribbean* clades are paraphyletic, and individuals from these previously defined populations are dispersed into multiple clades in the mitogenome tree (Figure 3). First, the clade *Mito 1* harbors 16 isolates in which 13 concordantly belong to the Arizona population using nucDNA genomes. The isolates B10813, Tucson2 and Tucson20 (*AZ clade I*) have conflicting phylogenetic distributions, previously these placed within *Texas/Mexico/SouthAmerica* and *AZ clade I* (Figure 3). Second, the clade *Mito II* contains the Venezuela group and three strains from the Arizona population (Tucson3, Tucson4 and Tucson14), although the Venezuela mitochondrial and nucDNA topologies are congruent. The next clade of *C. posadasii* is composed of four strains from Guatemala, and is perfectly concordant with the nucDNA tree. The clade *Mito III* is composed of two isolates from the *AZ Clade I* (Tucson6 and Tucson16) and two others from the *Caribbean* clade, and this lineage also shows conflicting phylogenetic placement (Figure 3). Finally, the clade *Mito IV* is composed primarily of isolates from the *Texas/Mexico/South America* clade.

Both nucDNA and mtDNA phylogenies revealed a clade composed of strains from Washington (16) which is genetically distinct from the rest of *C. immitis* (Figure 3). No other consistent pattern of clustering was observed for the remaining *C. immitis* individuals comparing the two phylogenies (Figure 3).

In general, our results suggest that nucDNA and mtDNA have similar evolutionary trajectories with no evidence of interspecific mtDNA exchange, but also the existence of phylogenetic incongruence at recent scales possibly due to within-species recombination.

## Discussion

Analysis of mitochondrial genomes as molecular markers confirms that the *Coccidioides* genus contains two species: *C. immitis* and *C. posadasii*. Mitochondrial markers are extensively used as molecular markers in speciation studies, including for *Coccidioides* (19, 37, 38). However molecular systematics of *Coccidioides* based on mitochondrial genes may lead to ambiguous conclusions at an intraspecies population level. Interestingly, both the mtDNA phylogeny and admixture plots of two distinct and divergent populations *C. immitis* Washington and *C. posadasii* Venezuela clearly reveal monophyly, as they are both reciprocally homozygous and no mixed genotypes are found (Figure 3). Both of these populations have emerged within the last 6,000 years according to estimates (13, 16). This suggests a strong founder effect followed by asexual reproduction in two endemic areas of the disease (39).

The evolutionary trajectories of both mtDNA and nucDNA genomes have been investigated using next generation sequencing data. Certainly, mtDNA and nucDNA genotypic incompatibilities may exist, as well as undetermined effects of cross-species hybridization and introgression (40). This is due in part to the fact that mitochondrial replication and division is not synchronized with nuclear division, and cells can contain numerous mitochondria, which may not undergo genetic recombination and may increase in number without cell division (41). Conflicting phylogenetic and population distributions have been observed in other pathogenic fungi, and our results indicate shared ancestry among recently diverged *C. immitis* and *C. posadasii* populations. For example, *Paracoccidioides brasiliensis* and *P. restrepiensis* appear to be polyphyletic using mtDNA markers, and the tree topologies differ from those obtained from nucDNA markers (42). Moreover, it is suggested that mitochondrial interspecific hybridization and introgression occurs in *Paracoccidioides* (42). Mitochondrial genomes can be difficult to assemble if high heterozygosity exists, as observed for the opportunistic pathogen *Candida metapsilosis*, which is part of the *C. parapsilosis* complex (43). This novel pathogen is a result of a hybridization event, which was detected in part by analyzing the mitochondrial genome. Within the primary pathogen *Cryptococcus gattii* complex, incongruences between mitochondrial and nuclear genes have been also reported (8). Mitochondrial genotypic and consequent phenotypic variation among these pathogen complexes is associated with virulence traits (44). These patterns are not restricted to human fungal pathogens. Some strains of the plant fungal pathogen *Verticillium longisporum* present a mosaic mitochondrial genome structure due to bi-parental inheritance impacting niche adaptation (45). Population genomic analyses of the lichen-forming fungi *Rhizoplaca melanophthalma* species complex suggest that hybridization and recombination in mitochondria might play a role in the speciation process of these symbiotic fungi (46).

The pathogenic lifestyles of *C. immitis* and *C. posadasii* necessitate potentially endozoan lifestyles (47) leading us to hypothesize that virulence, thermo-adaptation and oxidative stress could be driving genetic differentiation in mtDNA in *Coccidioides* species and populations. For example, in *Saccharomyces*, specific mutations the *cox1* gene in the mtDNA are associated with adaptation to variable temperatures. The authors suggest that the yeast mitochondrial genome is a hotspot in the evolution of thermal adaptation in *Saccharomyces* species (48, 49). *C. posadasii* is more heat tolerant than *C. immitis* and private alleles found in the mitochondrial genome might be responsible for this interspecific phenotypic variation. Importantly, in this manuscript we provided high-quality assemblies and annotations for the *Coccidioides* mitogenomes, which will facilitate deeper investigations into the impact of mitochondrial evolution in *Coccidioides’* niche adaptation, with particular emphasis on mammalian host co-evolution and oxidative stress responses.

## Author Contributions

The manuscript was written and edited by MT, JES, BFL, DRM and BMB. Data was analyzed by MT and BFL. Funding provided by BMB.

## Acknowledgments

We thank The Translational Genomics Institute and the Pathogen and Microbiome Institute and Northern Arizona University computing resources for data processing support. Special thanks to Dr. John Taylor for helpful discussion and edits.

## Supplementary Material

Accessions and details are listed in **Table S1**.

## References

1. Liu W, Cai Y, Zhang Q, Chen L, Shu F, Ma X, Bian Y. 2020. The mitochondrial genome of Morchella importuna (272.2 kb) is the largest among fungi and contains numerous introns, mitochondrial non-conserved open reading frames and repetitive sequences. Int J Biol Macromol 143:373–381.

2. Bullerwell CE, Leigh J, Forget L, Lang BF. 2003. A comparison of three fission yeast mitochondrial genomes. Nucleic Acids Res 31:759–68.

3. Aguileta G, de Vienne DM, Ross ON, Hood ME, Giraud T, Petit E, Gabaldon T. 2014. High variability of mitochondrial gene order among fungi. Genome Biol Evol 6:451–65.

4. Pogoda CS, Keepers KG, Nadiadi AY, Bailey DW, Lendemer JC, Tripp EA, Kane NC. 2019. Genome streamlining via complete loss of introns has occurred multiple times in lichenized fungal mitochondria. Ecol Evol 9:4245–4263.

5. Zubaer A, Wai A, Hausner G. 2018. The mitochondrial genome of Endoconidiophora resinifera is intron rich. Sci Rep 8:17591.

6. Lang BF, Laforest MJ, Burger G. 2007. Mitochondrial introns: a critical view. Trends Genet 23:119–25.

7. Stoddard BL. 2011. Homing endonucleases: from microbial genetic invaders to reagents for targeted DNA modification. Structure 19:7–15.

8. Bovers M, Hagen F, Kuramae EE, Boekhout T. 2009. Promiscuous mitochondria in Cryptococcus gattii. FEMS Yeast Research 9:489–503.

9. Barker BM, Litvintseva AP, Riquelme M, Vargas-Gastelum L. 2019. Coccidioides ecology and genomics. Med Mycol 57:S21–S29.

10. Lacy GH, Swatek FE. 1974. Soil ecology of Coccidioides immitis at Amerindian middens in California. Appl Microbiol 27:379–88.

11. Kollath DR, Miller KJ, Barker BM. 2019. The mysterious desert dwellers: Coccidioides immitis and Coccidioides posadasii, causative fungal agents of coccidioidomycosis. Virulence 10:222–233.

12. Fisher MC, Koenig GL, White TJ, San-Blas G, Negroni R, Alvarez IG, Wanke B, Taylor JW. 2001. Biogeographic range expansion into South America by Coccidioides immitis mirrors New World patterns of human migration. Proc Natl Acad Sci U S A 98:4558–62.

13. Teixeira MM, Alvarado P, Roe CC, Thompson GR, 3rd, Patane JSL, Sahl JW, Keim P, Galgiani JN, Litvintseva AP, Matute DR, Barker BM. 2019. Population Structure and Genetic Diversity among Isolates of Coccidioides posadasii in Venezuela and Surrounding Regions. mBio 10.

14. Teixeira MM, Barker BM, Stajich JE. 2019. Improved Reference Genome Sequence of Coccidioides immitis Strain WA_211, Isolated in Washington State. Microbiol Resour Announc 8.

15. McCotter OZ, Benedict K, Engelthaler DM, Komatsu K, Lucas KD, Mohle-Boetani JC, Oltean H, Vugia D, Chiller TM, Sondermeyer Cooksey GL, Nguyen A, Roe CC, Wheeler C, Sunenshine R. 2019. Update on the Epidemiology of coccidioidomycosis in the United States. Med Mycol 57:S30–S40.

16. Engelthaler DM, Roe CC, Hepp CM, Teixeira M, Driebe EM, Schupp JM, Gade L, Waddell V, Komatsu K, Arathoon E, Logemann H, Thompson GR, 3rd, Chiller T, Barker B, Keim P, Litvintseva AP. 2016. Local Population Structure and Patterns of Western Hemisphere Dispersal for Coccidioides spp., the Fungal Cause of Valley Fever. MBio 7:e00550–16.

17. Maxwell CS, Mattox K, Turissini DA, Teixeira MM, Barker BM, Matute DR. 2019. Gene exchange between two divergent species of the fungal human pathogen, Coccidioides. Evolution 73:42–58.

18. Neafsey DE, Barker BM, Sharpton TJ, Stajich JE, Park DJ, Whiston E, Hung CY, McMahan C, White J, Sykes S, Heiman D, Young S, Zeng Q, Abouelleil A, Aftuck L, Bessette D, Brown A, FitzGerald M, Lui A, Macdonald JP, Priest M, Orbach MJ, Galgiani JN, Kirkland TN, Cole GT, Birren BW, Henn MR, Taylor JW, Rounsley SD. 2010. Population genomic sequencing of *Coccidioides* fungi reveals recent hybridization and transposon control. Genome research 20:938–46.

19. Hamm PS, Hutchison MI, Leonard P, Melman S, Natvig DO. 2019. First Analysis of Human Coccidioides Isolates from New Mexico and the Southwest Four Corners Region: Implications for the Distributions of C. posadasii and C. immitis and Human Groups at Risk. J Fungi (Basel) 5.

20. Bankevich A, Nurk S, Antipov D, Gurevich AA, Dvorkin M, Kulikov AS, Lesin VM, Nikolenko SI, Pham S, Prjibelski AD, Pyshkin AV, Sirotkin AV, Vyahhi N, Tesler G, Alekseyev MA, Pevzner PA. 2012. SPAdes: a new genome assembly algorithm and its applications to single-cell sequencing. J Comput Biol 19:455–77.

21. Langmead B, Salzberg SL. 2012. Fast gapped-read alignment with Bowtie 2. Nat Methods 9:357–9.

22. Quinlan AR. 2014. BEDTools: The Swiss-Army Tool for Genome Feature Analysis. Current Protocols in Bioinformatics 47:11.12.1–11.12.34.

23. Cardoso MA, Tambor JH, Nobrega FG. 2007. The mitochondrial genome from the thermal dimorphic fungus Paracoccidioides brasiliensis. Yeast 24:607–16.

24. Tambor JH, Guedes RF, Nobrega MP, Nobrega FG. 2006. The complete DNA sequence of the mitochondrial genome of the dermatophyte fungus Epidermophyton floccosum. Curr Genet 49:302–8.

25. Wu Y, Yang J, Yang F, Liu T, Leng W, Chu Y, Jin Q. 2009. Recent dermatophyte divergence revealed by comparative and phylogenetic analysis of mitochondrial genomes. BMC Genomics 10:238.

26. Shang Y, Xiao G, Zheng P, Cen K, Zhan S, Wang C. 2016. Divergent and Convergent Evolution of Fungal Pathogenicity. Genome Biol Evol 8:1374–87.

27. Whiston E, Zhang Wise H, Sharpton TJ, Jui G, Cole GT, Taylor JW. 2012. Comparative transcriptomics of the saprobic and parasitic growth phases in Coccidioides spp. PloS one 7:e41034.

28. Greiner S, Lehwark P, Bock R. 2019. OrganellarGenomeDRAW (OGDRAW) version 1.3.1: expanded toolkit for the graphical visualization of organellar genomes. Nucleic Acids Res 47:W59–W64.

29. Li H, Durbin R. 2009. Fast and accurate short read alignment with Burrows-Wheeler transform. Bioinformatics 25:1754–60.

30. McKenna A, Hanna M, Banks E, Sivachenko A, Cibulskis K, Kernytsky A, Garimella K, Altshuler D, Gabriel S, Daly M, DePristo MA. 2010. The Genome Analysis Toolkit: a MapReduce framework for analyzing next-generation DNA sequencing data. Genome Res 20:1297–303.

31. Kurtz S, Phillippy A, Delcher AL, Smoot M, Shumway M, Antonescu C, Salzberg SL. 2004. Versatile and open software for comparing large genomes. Genome Biol 5:R12.

32. Nguyen LT, Schmidt HA, von Haeseler A, Minh BQ. 2015. IQ-TREE: a fast and effective stochastic algorithm for estimating maximum-likelihood phylogenies. Mol Biol Evol 32:268–74.

33. Kalyaanamoorthy S, Minh BQ, Wong TKF, von Haeseler A, Jermiin LS. 2017. ModelFinder: fast model selection for accurate phylogenetic estimates. Nat Methods 14:587–589.

34. Minh BQ, Nguyen MA, von Haeseler A. 2013. Ultrafast approximation for phylogenetic bootstrap. Mol Biol Evol 30:1188–95.

35. Puigbò P, Garcia-Vallvé S, McInerney JO. 2007. TOPD/FMTS: a new software to compare phylogenetic trees. Bioinformatics 23:1556–1558.

36. Van Dyke MCC, Teixeira MM, Barker BM. 2019. Fantastic yeasts and where to find them: the hidden diversity of dimorphic fungal pathogens. Curr Opin Microbiol 52:55–63.

37. Schroder MS, Martinez de San Vicente K, Prandini TH, Hammel S, Higgins DG, Bagagli E, Wolfe KH, Butler G. 2016. Multiple Origins of the Pathogenic Yeast Candida orthopsilosis by Separate Hybridizations between Two Parental Species. PLoS Genet 12:e1006404.

38. van de Sande WW. 2012. Phylogenetic analysis of the complete mitochondrial genome of Madurella mycetomatis confirms its taxonomic position within the order Sordariales. PLoS One 7:e38654.

39. Ballard JW, Whitlock MC. 2004. The incomplete natural history of mitochondria. Mol Ecol 13:729–44.

40. Sota T, Vogler AP. 2001. Incongruence of mitochondrial and nuclear gene trees in the Carabid beetles Ohomopterus. Syst Biol 50:39–59.

41. Mendoza H, Perlin MH, Schirawski J. 2020. Mitochondrial Inheritance in Phytopathogenic Fungi-Everything Is Known, or Is It? Int J Mol Sci 21.

42. Turissini DA, Gomez OM, Teixeira MM, McEwen JG, Matute DR. 2017. Species boundaries in the human pathogen Paracoccidioides. Fungal Genet Biol 106:9–25.

43. Pryszcz LP, Nemeth T, Saus E, Ksiezopolska E, Hegedusova E, Nosek J, Wolfe KH, Gacser A, Gabaldon T. 2015. The Genomic Aftermath of Hybridization in the Opportunistic Pathogen Candida metapsilosis. PLoS Genet 11:e1005626.

44. Ma H, May RC. 2010. Mitochondria and the regulation of hypervirulence in the fatal fungal outbreak on Vancouver Island. Virulence 1:197–201.

45. Depotter JRL, Beveren Fv, C.M. van den Berg G, Wood TA, Thomma BPHJ, Seidl MF. 2018. Nuclear and mitochondrial genomes of the hybrid fungal plant pathogen <em>Verticillium longisporum</em> display a mosaic structure. bioRxiv doi: 10.1101/249565:249565.

46. Keuler R, Garretson A, Saunders T, Erickson RJ, St. Andre N, Grewe F, Smith H, Lumbsch HT, Huang J-P, St. Clair LL, Leavitt SD. 2020. Genome-scale data reveal the role of hybridization in lichen-forming fungi. Scientific Reports 10:1497.

47. Taylor JW, Barker BM. 2019. The endozoan, small-mammal reservoir hypothesis and the life cycle of Coccidioides species. Med Mycol 57:S16–S20.

48. Li XC, Peris D, Hittinger CT, Sia EA, Fay JC. 2019. Mitochondria-encoded genes contribute to evolution of heat and cold tolerance in yeast. Sci Adv 5:eaav1848.

49. Baker EP, Peris D, Moriarty RV, Li XC, Fay JC, Hittinger CT. 2019. Mitochondrial DNA and temperature tolerance in lager yeasts. Sci Adv 5:eaav1869.

